# A Multi-Omics Processing Pipeline (MOPP) for Extracting Taxonomic and Functional Insights from Metaribosome Profiling (metaRibo-Seq) data

**DOI:** 10.64898/2026.03.11.710980

**Authors:** Yuhan Weng, Oriane Moyne, Corinn Walker, Eli Haddad, Chloe Lieng, Loryn Chin, Gibraan Rahman, Daniel McDonald, Rob Knight, Karsten Zengler

## Abstract

Metaribosome profiling (metaRibo-Seq) enables genome-wide measurement of translation across complex microbial communities by sequencing ribosome-protected mRNA fragments, but the short length of these footprints creates substantial nonspecific mapping against large reference genome collections, leading to spurious taxonomic and functional assignments. Here we present MOPP (Multi-Omics Processing Pipeline), a modular reference-based workflow that denoises metaRibo-Seq data by leveraging matched metagenomic coverage breadth to identify genomes likely to be truly present in a sample before aligning metatranslatomic and optional metatranscriptomic reads. MOPP generates taxon-by-gene count tables across genomic, transcriptional and translational layers, enabling integrated downstream analyses of microbial function. We evaluated MOPP using a defined 79-member synthetic human gut community profiled by metagenomics and metaRibo-Seq. Coverage breadth filtering markedly improved detection accuracy relative to a standard baseline workflow, with performance remaining robust across a broad intermediate threshold range and peaking at 92-95% coverage breadth. At a 92% threshold, MOPP reduced the number of distinct detected operational genomic units by 99.4% while retaining 87.8% of aligned metaRibo-Seq reads on average, and increased the F1 score from 0.02 to 0.61. Residual false positives were predominantly attributable to genomes with extremely high nucleotide similarity to true community members, whereas false negatives were enriched among low-abundance taxa, indicating that remaining errors are driven primarily by biological similarity and detection limits rather than widespread nonspecific mapping. Together, these results establish MOPP as a high-throughput workflow for robust processing of metaRibo-Seq in the context of matched metagenomics and position it as a scalable framework for integrated taxonomic and functional analysis of microbial communities across genomic, transcriptional and translational layers.

## INTRODUCTION

Metaribosome Profiling, or MetaRibo-Seq, is an emerging technique that quantifies genome-wide translational activities across microbes within complex communities by sequencing ribosome-protected mRNA fragments, commonly referred to as ribosome footprints^1^. By capturing actively translated genes, metaRibo-seq provides a direct readout of protein synthesis, bridging the gap between transcriptional potential and proteomic output.. Consequently, this approach reduces the interpretational discordance between transcript and protein levels that can limit metatranscriptomic approaches and enables the examination of microbial proteome allocation at a finer functional resolution^1,2^. As a sequencing-based translatomic readout, it naturally complements spectra-based metaproteomics and supports a more integrated multi-omic view, particularly where proteomic coverage and throughput remain limiting^2^.

Early applications of metaRibo-Seq have demonstrated its potential in uncovering translation of small proteins and peptides^2^ and in informing rational design of targeted microbiome interventions^1^. Despite its promise, metaRibo-seq remains in its early stages of development, and no standardized reference-based processing pipeline has yet been established.

One major analytical challenge in metaRibo-seq processing arises from the short length of ribosome footprints, typically 28–32 nucleotides in length^3,4^(**Fig. 1a**). These short reads often map identically and nonspecifically to multiple microbial genomes (**Fig. 1b**). This nonspecific mapping generates numerous spurious hits and complicates the interpretation of translational activities within the microbial communities^5^. One proposed strategy to address this limitation is *de novo* assembly of matched metagenomic reads followed by alignment of metaRibo-seq reads to the reconstructed assembly^4^. While this assembly-based approach can reduce misalignment to reference genomes that are distantly related to organisms present in the sample, it has been noted to be resource-intensive and often results in incomplete reference metagenomes^5^. Therefore, enabling a reference-based approach for rapid identification and characterization of metaRibo-Seq at scale requires alternative strategies to mitigate spurious read alignment.

**Fig. 1:**
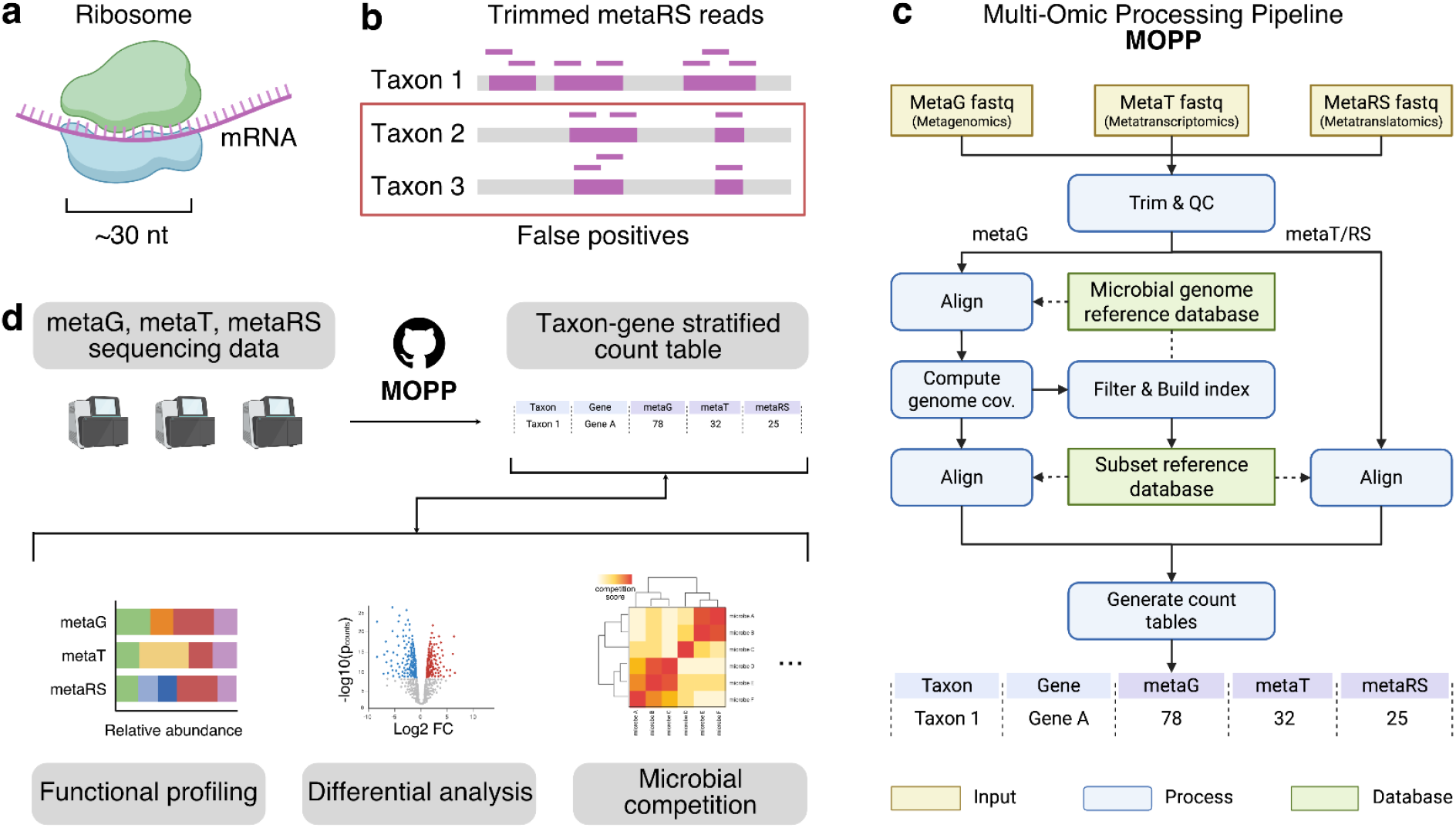
Overview of metaRibo-seq ambiguity and the MOPP workflow. a) Ribosome profiling captures ∼30 nt ribosome-protected mRNA fragments. b) After trimming, direct alignment of short metaRibo-seq reads to a general microbial genome reference database results in multi-alignment and false positives. c) Schematic of the Multi-Omics Processing Pipeline (MOPP), which integrates metagenomic (metaG), metatranscriptomic (metaT), and metatranslatomic (metaRS) data through quality control, alignment, genome indexing, and count table generation. d) MOPP produces taxon- and gene-resolved count tables that enable downstream analyses, including functional profiling, differential analysis, and prediction of microbial competition. Icons are created using BioRender. Knight Lab (2024). https://BioRender.com/.

Here, we propose a workflow that leverages genome coverage breadth calculated from matched metagenomic reads to reduce spurious metaRibo-seq alignments (**Fig. 1c**). Coverage breadth, defined as the proportion of a reference genome covered by at least one sequencing read^6,7^, has been shown to outperform conventional filtering metrics such as relative abundance and prevalence in reducing false positives^8,9^. For organisms present in a sample, metagenomic coverage breadth can approach 100% when a large number of samples containing them have been sequenced. In contrast, this assumption does not hold for metatranscriptomic or metatranslatomic reads, which capture only transcribed or translated regions rather than the full genome. Consequently, coverage breadth cannot be directly applied to RNA-seq or metaRibo-seq to distinguish truly present organisms from potential false positives. Building upon this distinction, our approach uses metagenomic coverage breadth to identify candidate reference genomes prior to aligning metaRibo-seq reads, thereby reducing multimapping and minimizing alignment to distantly related genomes absent from the sample.

To implement this strategy, we developed Multi-Omics Processing Pipeline (MOPP), a reference-based workflow designed to process metaRibo-seq data in the context of matched metagenomic sequencing. MOPP optionally integrates matched metatranscriptomic data within the same analytical framework. The primary output of MOPP is a taxon-by-gene stratified count table spanning genomic, transcriptional, and translational layers. These multi-omic matrices enable downstream analyses including functional profiling, differential expression analysis, and inference of microbial competition dynamics^1^ (**Fig. 1d**).

Throughout this study, we refer to metagenomic, metatranscriptomic, and metatranslatomic sequencing as metaG, metaT, and metaRS, respectively. MetaRS and metaRibo-Seq are used interchangeably.

## RESULTS

### Overview of MOPP workflow

MOPP integrates matched metaG, metaT, and metaRibo-Seq data to generate a multi-omic feature table detailing per-taxon, per-gene counts for each sample. The workflow begins with trimming and quality control of all sequencing reads. Trimmed DNA-Seq reads are then aligned to the Web of Life (WoL)^10^ database by default, or a user-specified reference database, and coverage breadth is calculated for all aligned reference genomes. Genomes with coverage breadth higher than a user-specified threshold are extracted to construct a custom subset reference database retaining only the organisms that are well-represented in the sample and thus likely to be truly present. MetaRibo-Seq and RNA-Seq reads previously trimmed with the same process are then aligned to this subset database to produce the final taxon-gene stratified table. A log file is generated to record the start and stop times of each step, aiding in performance tracking and debugging (**Fig. 1c**; Methods).

MOPP is highly modular. Users can invoke individual steps (e.g. trimming, coverage breadth calculation .etc) via standalone commands (e.g. mopp trim, mopp cov .etc), allowing integration with external tools and convenient re-execution with different parameters. Additionally, MOPP is flexible, allowing users to incorporate their own reference database for studies focused on specific genomes of interest.

### Coverage-based genome filtering improves detection accuracy

We evaluated the performance of MOPP using a defined synthetic microbial community (SynCom). A 79-member synthetic human gut microbial community profiled by metaG and metaRibo-Seq was used as a benchmarking dataset (**Fig. 2a**; Methods). Two key metrics, the number of distinct operational genomic units (OGUs) and the number of aligned reads, were recorded and compared between the standard sequencing processing pipeline (baseline) and MOPP (Methods). A substantial reduction in the number of distinct OGUs, accompanied by minimal loss of aligned reads, would indicate that MOPP effectively denoises metaRibo-Seq data. Since the true community composition was known, this system also enabled direct evaluation of F1 score, which reflects both precision (the fraction of detected organisms that are truly present) and recall (the fraction of true organisms that are detected), providing a balanced measure of detection accuracy (Methods).

**Fig. 2:**
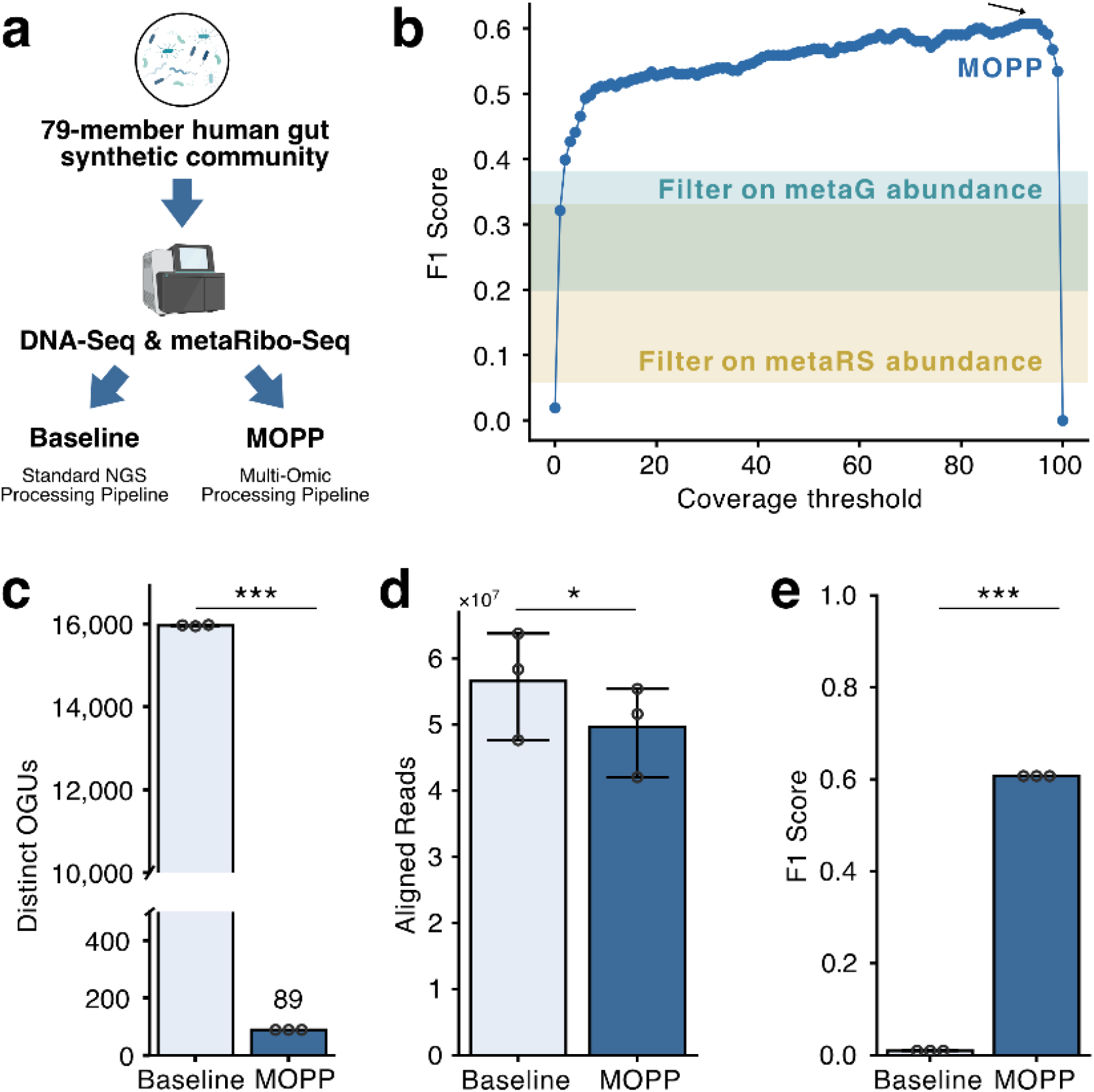
MOPP denoises metaRibo-seq while preserving read depth. a) Experimental design using an 80-member synthetic human gut community profiled by metaG and metaRibo-seq and processed with or without MOPP. b) F1 score versus coverage threshold. Arrow points to the maximum F1 score of 0.60 at coverage breadth threshold 92-95%. Teal shaded area: the range of F1 scores achieved by metagenomic relative abundance filtering using common thresholds 0.001%-1%. Yellow shaded area: the range of F1 scores achieved by metatranslatomic relative abundance filtering using common thresholds 0.001%-1%. c) MOPP reduces the number of distinct Operational Genomic Units (OGUs) detected by 99.4% at the coverage threshold of 92% (Paired t-test, *t* = -1634.18, *p* = 3.74×10^-7^). d) Total aligned reads are largely preserved, with on average a 87.8% retention at a 92% coverage threshold after MOPP processing. (Paired t-test, *t* = -8.62, df = 2, *p* = 1.32×10^-2^). e) F1 scores between baseline and MOPP at a 92% coverage threshold. MOPP increased F1 from 0.02 to 0.61 (Paired t-test, *t* = 9.88×10^4^, *p* = 1.02×10^-10^).

We first observed that increasing the coverage breadth threshold rapidly improves F1 score at low cutoffs, after which performance enters a broad plateau: F1 varies only modestly across intermediate thresholds (∼10–95% coverage). A maximum F1 score of 0.60 is achieved between 92-95% coverage, whereas more stringent thresholds lead to pronounced performance drops (**Fig. 2b**). This pattern is consistent with the empirical distribution of metagenomic coverage breadth. The majority of detected organisms exhibit very low coverage breadth (<10%; **Suppl. Fig. 1a**), suggesting the presence of high false-positive hits, possibly driven by sequence similarity and nonspecific mapping. In contrast, taxa that are truly present are enriched at high coverage breadth, peaking around ∼80-100% (**Suppl. Fig. 1a**). Together, these distributions explain why moderate breadth thresholds improve performance by removing low-coverage false positives, while overly stringent cutoffs can exclude true community members that fall just below the threshold due to low abundance and/or incomplete metagenomic coverage, thereby increasing false negatives and reducing overall detection performance. Notably, coverage-based filtering attains higher F1 scores than those achievable by relative-abundance filtering across commonly used thresholds (0.001%-1%), whether applied directly to metaRibo-Seq or derived from paired metagenomic abundance (**Fig. 2b**).

At a 92% coverage threshold, MOPP substantially reduced the number of distinct OGUs while retaining most aligned reads. The baseline processing workflow reported 15,942 to 15,974 distinct OGUs across triplicates, despite only 79 taxa being truly present. In contrast, under the MOPP workflow, each sample retained 89 OGUs, a 99.4% reduction of the number of distinct taxa reported (**Fig. 2c, Suppl. Table 1**). This reduction was consistent across all three paired samples, and paired t-test on the within-sample differences between pipelines indicated a large and statistically significant decrease in distinct OGUs (Paired t-test, *t* = -1634.18, *p* = 3.74×10^-7^). With only three paired samples, statistical inference is limited; we therefore emphasize the effect size and consistency of results across matched pairs rather than relying on *p*-values alone.

Despite substantial decrease in the number of distinct OGUs, read retention remained high. The number of aligned metaRibo-Seq reads decreased from 47.7-63.8 million reads under baseline processing to 42.1-55.4 million reads after MOPP, corresponding to 87.8% read retention on average (**Fig. 2d**). The paired t-test likewise supported a significant but comparatively smaller reduction in aligned reads (Paired t-test, *t* = -8.62, df = 2, *p* = 1.32×10^-2^).

Consistent with the large reduction in OGUs and high retention in aligned reads, MOPP improved F1 score-based detection accuracy. F1 scores were near zero across samples (0.02 on average) in the baseline, whereas MOPP increased F1 to 0.61 for all three samples (a 59.9% increase on average) using MOPP (**Fig. 2e**). The improvement was highly consistent across pairs, and the paired t-test on per-sample differences confirmed a significant increase in F1 (*t* = 9.88×10^4^, *p* = 1.02×10^-10^).

Given the small number of paired samples (*n* = 3), we interpret *p*-values cautiously and focus primarily on the magnitude and consistency of effects across matched pairs. We also applied the Wilcoxon signed-rank test to each metric; with *n* = 3 and all paired differences sharing the same sign, the exact two-sided Wilcoxon test yields a coarse *p*-value of 0.25 and is therefore underpowered to distinguish strong effects in this setting. We therefore report Wilcoxon results as a robustness check while using the paired *t*-test to summarize mean paired differences. While one of the best-performing coverage breadth thresholds, 92%, is chosen for detailed analyses here, we note that mid-range thresholds (∼10–95%) do not strongly affect the number of distinct OGUs (**Suppl. Fig. 1b**) and error rates (**Suppl. Fig. 1c,d**) and the MOPP performance is robust to the selection of thresholds in this range (**Suppl. Fig. 2a-d**).

Overall, these results demonstrate that coverage breadth-based genome filtering with MOPP markedly improves detection accuracy by removing low-breadth false positives while preserving the vast majority of aligned reads, yielding a denoised OGU profile that closely recapitulates the known synthetic community composition.

### False positives are driven by high genomic similarity

Because the true community composition is known, we quantified false positives and false negatives directly using confusion matrices. Baseline processing produced pervasive false-positive detections, with thousands of taxa spuriously identified despite successful recovery of true community members (**Fig. 3a**). In contrast, applying MOPP markedly reduced false positives while retaining most true positives (**Fig. 3b**). At a 92% coverage-breadth threshold, MOPP substantially outperforms the baseline across all performance metrics (Table 1). Although the baseline achieves perfect recall (1.00), it does so with extremely poor precision (0.00971) and low specificity (0.495), resulting in near-chance accuracy (0.498) and a very low F1 score (0.0192), consistent with pervasive false-positive calling. In contrast, MOPP maintains strong recall (0.646) while dramatically improving precision (0.573) and specificity (0.998), yielding high overall accuracy (0.996) and an F1 score of 0.607. Notably, F1 provides a balanced metric between precision and recall, and the large increase in F1 (0.0192 to 0.607) indicates that MOPP achieves a far more favorable trade-off between recovering true positives and limiting false positives than the baseline approach (**Table 1**).

**Fig. 3:**
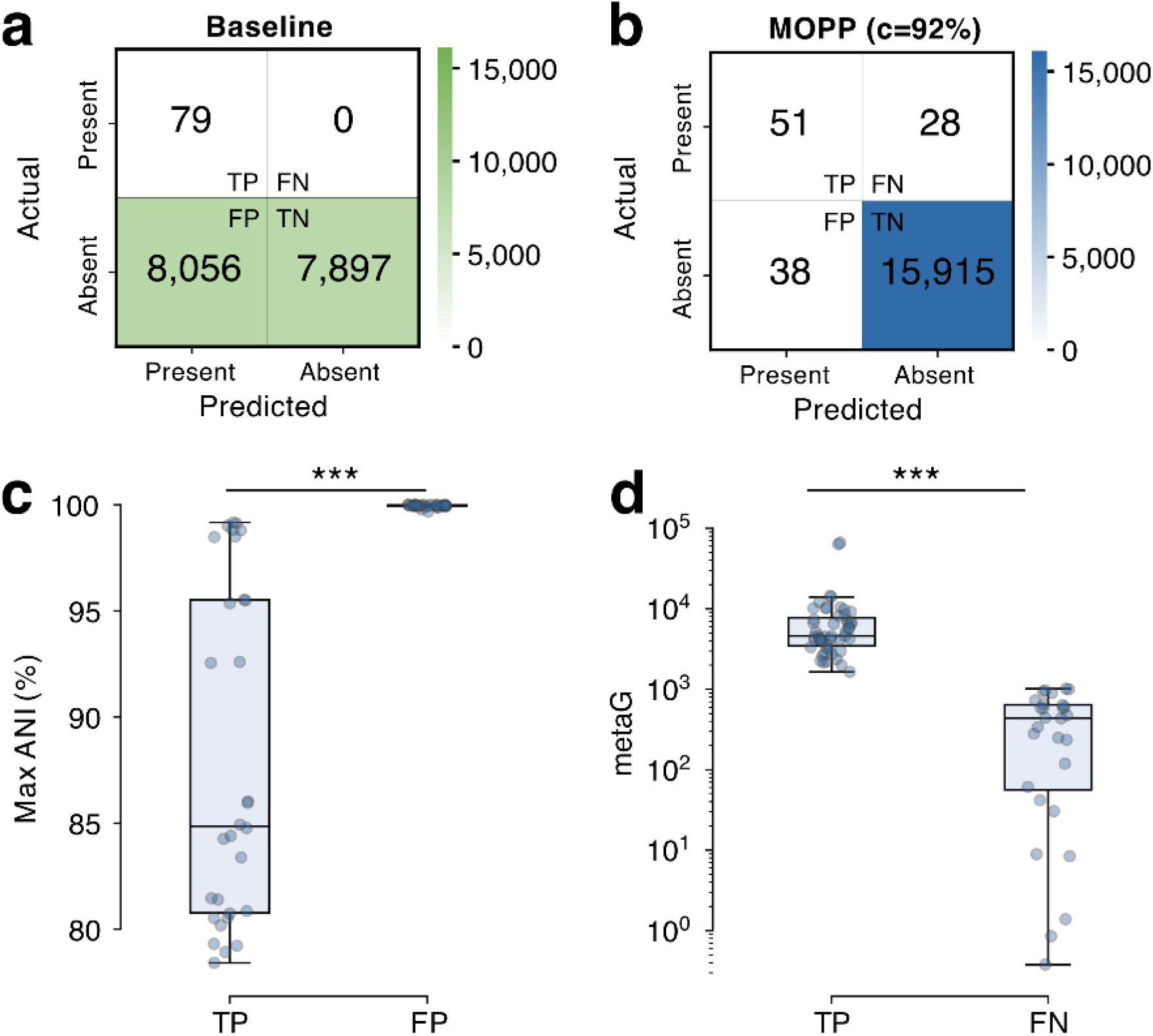
Characterization of false positives and false negatives in metaRibo-seq detection. a) Confusion matrix for baseline processing, revealing extensive (8,056) false-positive detections despite correct identification of true community members. Top left: true positive (TP), top right: false negative (FN), bottom left: false positive (FP), bottom right: true negative (TN). b) Confusion matrix after MOPP processing with a 92% genome coverage threshold, showing substantial reduction of false positives while retaining most true positives. Top left: TP, top right: FN, bottom left: FP, bottom right: TN. c) Maximum average nucleotide identity (ANI) of TPs and FPs to the most similar non-self reference genome among the 80-member community (for TPs) or to any community member (for FPs), showing that FPs are typically highly similar to at least one true genome, consistent with spurious alignments driven by closely related sequences (Mann-Whitney U test, two-sided, *p* = 5.12×10^-12^). d) b) Distribution of average metagenomic relative abundance (DNA-Seq) for TPs and FNs, showing that false negatives are significantly lower in abundance (Mann–Whitney U test, two-sided, *U* = 1,428, *p* = 2.62×10^-13^).

At a 92% coverage-breadth threshold, MOPP nevertheless yielded 38 false positives. We compared the maximum average nucleotide identity (ANI) of these false positives to the true community members. All of them have 100% ANI to at least one of the 79 organisms present in the SynCom (**Fig. 3c**). On average, these false positives exhibited significantly higher average nucleotide identity (ANI) to true community members than did the true positives themselves, excluding self-comparisons (Mann-Whitney U test, two-sided, *p* = 5.12×10^-12^; excluding comparisons with ANI much lower than 80% (Methods); **Fig. 3c, Suppl. Fig. 3a**). These results showed that metaRibo-Seq reads align to conserved coding regions shared among closely related genomes, producing residual false-positive detections even under stringent coverage-based filtering.

### False negatives are associated with low abundance

We next examined biological and technical factors associated with false-negative detections. False-negative taxa showed significantly lower metagenomic abundance relative to true positives (Mann–Whitney U test, two-sided, *U* = 1,428, *p* = 2.62×10^-13^; **Fig. 3d**).

Coverage analyses further indicated that true positive community members achieve increasing coverage with rising abundance, whereas spurious detections often display localized mapping patterns despite moderate read abundance (**Suppl. Fig. 3b**). These observations reinforce genome coverage breadth as a robust discriminator between genuine organism detection and off-target alignments.

Finally, although strains were individually grown to a common OD and mixed at equal ratios, growth rates varied substantially across the 80-member community. This increases the time and handling needed to synchronize cultures and may introduce variability (e.g., differences in physiological state, OD-to-biomass conversion, or viability) that could contribute to non-detection of some taxa.

In conclusion, we introduced MOPP, a reference-based workflow for processing metaRibo-Seq data in the context of metagenomics, and we validated MOPP using a defined synthetic microbial community and further evaluated its performance in human fecal microbiome samples spanning both infant and adult cohorts. These analyses demonstrate MOPP’s effectiveness in reducing spurious alignment of metaRibo-seq while preserving quantitative translational signal in both infant fecal samples and more complex adult fecal samples.

## METHODS

### MOPP Workflow

MOPP is a reference-based workflow for extracting taxonomic and functional information from metaRibo-Seq in microbial communities that are well represented by available reference genomes (e.g. defined SynComs and well-characterized environments such as human stool). It takes in matched DNA-Seq, RNA-Seq, and metaRibo-Seq data from the same sample(s) and generates a multi-omic feature table enumerating the gene and taxon counts in each sample across omic levels. Specifically, in the MOPP workflow, sequencing reads are first trimmed using TrimGalore (Cutadapt) version 1.1841^11^ with default trimming parameters and quality controlled using FastQC version 0.11.942^12^. Trimmed DNA-Seq reads are then aligned to the Web of Life database (WoL)^10^, or any user-specified database, using bowtie2 version 2.3.243^13^, followed by the calculation of coverage breadth of aligned reference genomes using micov v1.0.0^7^. Based on the distribution of coverage breadth, a user-defined sample-specific threshold is used to filter reference genomes. Genomes with coverage breadth higher than the chosen threshold will be extracted and indexed into a subset reference genome database. Previously trimmed metaRibo-Seq and RNA-Seq reads will then be aligned to this subset database and generate the final count table. RNA-Seq is aligned using default parameters whereas metaRibo-Seq is aligned using a stricter parameter ‘score-min “L,0,-0.01”’ to account for the shorter nature of ribosome-protected fragments. A log file recording the start and stop times of each step will be generated in the output directory for tracking the time performance and debugging if needed.

### Assembly of the synthetic community

Culturing and assembly were performed in an anaerobic chamber maintained at 5% H_2_, 20% CO_2_, and 75% N2. A total of 79 bacterial strains (**Suppl. Table 2**) were individually inoculated from glycerol stocks and cultured in 10mL filter-sterilized modified brain-heart infusion (BHI) broth (**Suppl. Table 3**) at 37°C for 4 days without shaking. Optical density at 600nm (OD600) was measured using a Cerillo Alto microplate reader.

Each culture was normalized by diluting in modified BHI broth to an OD600 of 0.02 after subtraction the media blank reading. The human gut synthetic community (hCom) was assembled by combining 10mL of each normalized culture to generate an equal mix stock. This equal mix stock was then diluted 1:100 in modified BHI broth to complete the final hCom inoculum.

The initial time point (0 h) was collected prior to incubation. For DNA- and RNA-seq, 5mL of the hCom culture was aliquoted in triplicate and harvested by centrifugation. Cell pellets were resuspended in DNA/RNA Shield. For metaRibo-seq, 25mL of the culture was aliquoted in triplicate, supplemented with 50uL chloramphenicol (CAM) per sample, and harvested by centrifugation. Cell pellets were resuspended in RNAlater. All samples were immediately frozen in liquid nitrogen and stored at -80°C.

### DNA-Seq and RNA-Seq sample preparation

DNA and RNA extraction was performed using the ZymoBIOMICS DNA/RNA miniprep kit (Zymo) in accordance with the manufacturer’s instructions. rRNA was removed from RNA samples using the QIAseq FastSelect-5S/16S/23S kit (Qiagen). DNA-seq libraries were prepared using the Nextera XT DNA Library kit (Illumina, Cat# FC-131-1096 and FC-131-2001) and RNA-seq libraries were prepared using the KAPA RNA HyperPrep kit (Roche). Sequencing was performed on an Illumina NovaSeq 6000, PE100 platform.

### MetaRibo-Seq sample preparation

MetaRibo-Seq was performed using the adapted protocol for axenic bacterial cultures. Bacterial cells were mechanically lysed using the OMNI Bead Ruptor Elite in a solution containing chloramphenicol and Guanosine-5’-[(β,γ)-imido]triphosphate (GMPPNP) to halt protein synthesis. Briefly, mechanical bacterial lysis was performed in a solution containing chloramphenicol and Guanosine-5’-[(β,γ)-imido]triphosphate (GMPPNP) to stop protein elongation. Resulting lysates were treated with MNase and DNase to degrade nucleic acids that were not protected by ribosomes. Monosome recovery was performed using RNeasy Mini spin size-exclusion columns (Qiagen) and RNA Clean & Concentrator-5 kit (Zymo). rRNA removal was performed using the QIAseq FastSelect-5S/16S/23S kit (Qiagen). MetaRibo-Seq libraries were prepared using the NEBNext Small RNA Library Prep set for Illumina, with modifications (see details in Suppl. Material 1). Amplification was followed in real time using SYBR-Green and stopped when reaching a plateau^1^.

### Sequencing

Library concentrations following library preparation were quantified using the Qubit dsDNA HS Assay kit on the QuBit 2.0 Fluorometer. Library quality and fragment size were assessed using a 4200 TapeStation System. Libraries were sequenced on an Illumina NovaSeq, PE100 platform. Libraries were sequenced to 10 million reads for metagenomic samples, 50 million reads for metatranscriptomic samples, and 100 million reads for metatranslatomic samples.

### Data processing

In the default processing workflow (“Baseline”), sequencing reads were trimmed using TrimGalore (Cutadapt) version 1.1841 with default trimming parameters and quality controlled using FastQC version 0.11.94211. Trimmed reads are then aligned to the reference database (in this case, SynCom genomes and Web of Life 2 (WoL2)) using bowtie2 version 2.3.24312, followed by count table generation using Woltka v0.1.7^14^ with default parameters, which reflects a standard NGS processing pipeline^15^. The default mode is designed to retain a broad set of candidate taxa/features with the expectation that downstream analyses will apply study-specific filtering (e.g. focusing on phenotype-associated taxa). See ‘MOPP workflow’ section for details on the MOPP processing approach.

### SynCom Validation

To validate the MOPP workflow, we utilized the known composition of the synthetic community. True Positives (TP) were defined as taxa included in the synthetic community that were correctly detected by the workflow. False Positives (FP) were defined as taxa not included in the synthetic community but incorrectly identified in the output. False Negatives (FN) refers to taxa included in the synthetic community that the workflow failed to detect, whereas True Negatives (TN) are taxa from the reference database (WoL) that were correctly excluded as they were not present in the synthetic community. Samples were triplicated. We evaluated workflow performance using the following metrics:

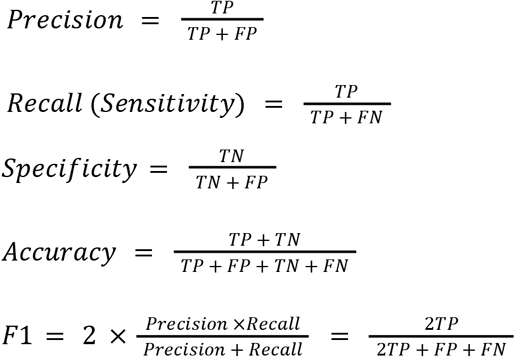

The F1-score is defined as the harmonic mean of precision and recall and was utilized as the primary metric to provide a balanced assessment of precision and recall.

Additionally, the distinct number of OGUs and number of aligned reads were evaluated. The distinct number of OGUs refers to the number of unique OGUs identified in each sample. Gram positive/negative status was assigned to each community member based on a literature review at the highest possible taxonomic resolution. To ensure accuracy and account for intra-generic variability, status was assigned at the strain level where possible, or at the species level as a minimum requirement; assignments were not inferred at the genus level or higher.

### False Positive and False Negative Analysis

ANIs were calculated using FastANI v1.32. In Figure 3c, the maximum ANI between each true positive detected and all 79 SynCom organisms were plotted in the left bar, excluding self-comparisons. Similarly, the maximum ANI between each false positive detected and all 79 SynCom organisms were plotted in the right bar. In Supplementary Figure 3a, instead of maximum ANIs, average ANIs were plotted. Comparisons with ANIs much lower than 80% were excluded as a built-in feature of FastANI.

## CODE AVAILABILITY

The MOPP workflow is available under an open-source license and can be obtained at https://github.com/sherlyn99/mopp.

## ACKNOWLEDGEMENTS

We are thankful to Dr. Michael Fischbach (Stanford University) for providing us with the strain for assembling the synthetic community. We thank Dr. Manish Kumar, Dr. Maxwell Neal, Dr. Rodrigo Santibáñez, and Dr. Juan Tibocha-Bonilla (UCSD) for their input on statistical analyses or/and manuscript.

## FUNDING

This material is based upon work supported by the U.S. Department of Energy, Office of Science, Office of Biological & Environmental Research under Awards DE-SC0021234 and DE-SC0022137. This work was also in part supported by the National Institute of Diabetes and Digestive and Kidney Diseases of the National Institutes of Health under award number 1R01DK140328-01. Furthermore, the development of the technologies described in this article were in part funded through Trial Ecosystem Advancement for Microbiome Science Program at Lawrence Berkeley National Laboratory funded by the U.S. Department of Energy, Office of Science, Office of Biological & Environmental Research Awards DE-AC02-05CH11231. The work was also supported by the UC San Diego Center for Microbiome Innovation (CMI) through a Grand Challenge Award and by a Seed grant made available through the UC San Diego Larsson-Rosenquist Foundation Mother-Milk-Infant Center of Research Excellence. The support of the Family Larsson-Rosenquist Foundation is gratefully acknowledged. This publication includes data generated at the UC San Diego IGM Genomics Center utilizing an Illumina NovaSeq 6000 that was purchased with funding from a National Institutes of Health SIG grant (#S10 OD026929).

## COMPETING INTERESTS

The authors declare the following competing interests: D.M. is a consultant for BiomeSense, Inc., has equity and receives income. R.K. is a scientific advisory board member, and consultant for BiomeSense, Inc., has equity and receives income. He is a scientific advisory board member and has equity in GenCirq. He has equity in and acts as a consultant for Cybele. K.Z. is co-founder of Isolation Bio, Native Microbials, and Guilden Corporation and has equity in all companies. He is also a co-founder and board member of the Prosper Soils Institute, a non-profit organization. The terms of these arrangements have been reviewed and approved by the University of California, San Diego in accordance with its conflict of interest policies. The remaining authors declare no competing interests.

## AUTHOR CONTRIBUTIONS

Y.W. conceived and designed the study. Y.W. and O.M. conceived the bioinformatic approach. C.W., C.L. and L.C. conducted experiments and collected data. Y.W. and E.H. developed the software package with the guidance from G.R. and D.M. on software development and micov usage. R.K. and K.Z. supervised the study. Y.W. performed the data processing and statistical analyses and wrote the manuscript with input from all co-authors.

**Supplementary Information is available for this paper**.

